# Cholecystokinin 1 Receptor (Cck1R) Normalizes mTORC1 signaling and is Protective to Purkinje cells of SCA Mice

**DOI:** 10.1101/2021.02.16.431490

**Authors:** Emily A.L. Wozniak, Zhao Chen, Sharan Paul, Praseuth Yang, Karla P. Figueroa, Jill Friedrich, Tyler Tschumperlin, Michael Berken, Melissa Ingram, Christine Henzler, Stefan M. Pulst, Harry T. Orr

**Author notes:** Authors contributed equally (Emily A.L. Wozniak, Zhao Chen, Sharan Paul).

## Abstract

Spinocerebellar Ataxias (SCAs) are a group of genetic diseases characterized by progressive ataxia and neurodegeneration, often in cerebellar Purkinje neurons. A SCA1 mouse model, *Pcp2-ATXN1[30Q]D776*, has severe ataxia in absence of progressive Purkinje neuron degeneration and death. Previous RNA-seq analyses identified cerebellar up-regulation of the peptide hormone Cholecystokinin (Cck) in *Pcp2-ATXN1[30Q]D776* mice. Importantly, absence of Cck1 receptor (Cck1R) in *Pcp2-ATXN1[30Q]D776* mice confers a progressive disease with Purkinje neuron death. A Cck1R agonist, A71623 administered to *Pcp2-ATXN1[30Q]D776;Cck^-/-^* and *Pcp2-AXTN1[82Q]* mice dampened Purkinje neuron pathology and associated deficits in motor performance. In addition, A71623 administration improved motor performance of *Pcp2-ATXN2[127Q]* SCA2 mice. Moreover, the Cck1R agonist A71623 corrected mTORC1 signaling and improved expression of calbindin in cerebella of *AXTN1[82Q]* and *ATXN2[127Q]* mice. These results indicate that manipulation of the Cck-Cck1R pathway is a potential therapeutic target for treatment of diseases involving Purkinje neuron degeneration.

## INTRODUCTION

Among inherited neurodegenerative diseases are the severe forms of the dominantly inherited spinocerebellar ataxia (SCA) that are due to an in-frame CAG trinucleotide expansion encoding a polyglutamine (polyQ) tract (Durr, 2010; Klockgether, 2011). These SCAs, SCA1, 2, 3, 6, 7, and 17, are clinically characterized by a progressive ataxia with cerebellar degeneration that in most of these SCAs consists of severe degeneration and loss of Purkinje cells, the major integrative neuron of the cerebellar cortex (Koeppen, 2005; Seidel et al., 2012).

To gain insight into molecular function of ATXN1 as well as cellular pathways critical for SCA1-like disease in Purkinje cells, a series of SCA1 mouse models were developed and characterized that manifest SCA1-like disease phenotypes including cerebellar ataxia and loss of Purkinje cells (Burright et al., 1995; Watase et al., 2002; Duvick et al., 2010). An outcome of this work is the concept that a critical ATXN1 function in relation to SCA1 Purkinje cell disease is its role in regulating transcription and RNA processing (Paulson et al., 2017). Two ATXN1 transgenic mouse lines formed the basis of a study that utilized RNA sequencing (RNA-seq) to reveal cerebellar gene expression changes associated with disease progression and protection (Ingram et al., 2016). In these mice, expression of ATXN1 is specifically directed to Purkinje cells. One line were mice expressing ATXN1 with an expanded polyQ, ATXN1[82Q], that manifest a progressive disease culminating in Purkinje cell death (Burright et al., 1995; Clark et al., 1997). The second line were mice expressing ATXN1 with a wild type polyQ stretch but with an Asp at position 776 that mimics some aspects of S776 phosphorylation. While this amino acid substitution transforms human wild type ATXN1[30Q] into a pathogenic protein with *ATXN1[30Q]-D776* mice manifesting ataxia that is as severe as manifested by *ATXN1[82Q]* animals. In contrast to *ATXN1[82Q]* mice, cerebellar disease in *ATXN1[30Q]-D776* mice do not manifest with a progressive cerebellar pathology that cumulates with Purkinje neuron death (Duvick et al., 2010). Analysis of the ATXN1 transgenic RNA-seq data demonstrated a substantial, unique and Purkinje cell-specific up regulation of cholecystokinin *(Cck)* RNA in *Pcp2-ATXN1[30Q]D776* mice (Ingram et al., 2016). Upon crossing *ATXN1[30Q]D776* mice to *Cck^-/-^* and *Cck1R^-/-^* mice, it was found that absence of either Cck or Cck1R in *Pcp2-ATXN1[30Q]D776* mice resulted in a Purkinje cell disease in which pathology progressed to cell death. These results strongly supported a hypothesis where elevated Cck and its subsequent processing to a peptide that binds to Purkinje cell Cck1R in an autocrine manner causes the lack of progressive Purkinje cell pathology in *ATXN1[30Q]D776* mice.

Here we show that administration of the Cck1R agonist, A71623, was protective against progressive ataxia and Purkinje cell pathology in *ATXN1[30Q]D776/ Cck^-/-^* . Moreover, RNA-seq analysis of *ATXN1[30Q]D776/ Cck^-/-^* cerebella disclosed a WGCNA gene module having a significant overlap with a gene network recently associated with Purkinje cell disease progression in ATXN1[82Q] mice (Ingram et al., 2016), indicating that the progressive disease in *ATXN1[30Q]D776/ Cck^-/-^* and *ATXN1[82Q]* mice have molecular features in common. Correspondingly, administration of the Cck1R agonist A71623 corrected mTORC1 signaling and was protective against progressive ataxia in *ATXN1[82Q*] mice. Lastly, administration of A71623 also improved and cerebellar mTORC1 signaling and motor performance in *ATXN2[127Q]* mice. These results provide proof of principle for Cck1R activation as a therapeutic option across the SCAs that are characterized by Purkinje cell dysfunction/degeneration.

## RESULTS

### Administration of a Cck1R Agonist A71623 Reduces Disease in *Pcp2-ATXN1[30Q]D776/ Cck^-/-^* Mice

*Pcp2-ATXN1[30Q]D776* animals have ataxia in absence of Purkinje cell progressive pathology (Duvick et al., 2010). Previously, we found that deletion of cholecystokinin (*Cck*) or cholecystokinin receptor 1 (*Cck1R)* genes from *ATXN1[30Q]D776* animals converted disease from being non-progressive to one in which Purkinje cell pathology is progressive and readily detected by 36 weeks of age (Figure S1). This lead to the suggestion that a Cck/Cck1R pathway protects *Pcp2-ATXN1[30Q]D776* Purkinje cells from a progressive pathology (Ingram et al., 2016). We reasoned that a test of this idea would be to assess whether activation of Purkinje cell Cck1Rs in *ATXN1[30Q]D776/ Cck^-/-^* mice dampened disease progression.

A71623, a Cck tetrapetide analogue, is highly selective agonist for Cck1R. In rodents, peripheral administration is able to elicit CNS-mediated behavioral effects (Asin et al., 1992a and 1992b). To assess the ability of A71623 administered peripherally to activate cerebellar Cck1Rs, phosphorylation of cerebellar Erk1 and Erk2, downstream targets of Cck1R activation (Dufresne et al., 2006), was assessed in mice following IP injections of A71623. Doses selected were 0.026mg (30nmoles)/Kg, a dose previously shown to have central effects in mice (Asin et al., 1992a), and 0.132mg/Kg the maximum dose given solubility of A71623. As shown in Figure S2, IP injections of A71623 at these doses resulted in a dose dependent increase in cerebellar Erk1 and Erk2 phosphorylation.

Figure 1A depicts the A71623 treatment strategy utilized to assess whether this Cck1R agonist impacts Purkinje cell disease progression in *ATXN1[30Q]D776/ Cck^-/-^* mice. Since the extent of recovery from ATXN1-Induced neurodegeneration decreases with age in *SCA1* transgenic mice (Zu et al., 2004), we selected to initiate Cck1R agonist treatment early in disease progression. Briefly, baseline motor assessments were performed at five- and six-weeks using balance beam and Rotarod, respectively. Upon completion of baseline Rotarod assessments, osmotic pumps were implanted intraperitoneally and the Cck1R agonist A71623 was administered at a rate of 0.026mg (30nmoles)/Kg/day. Balance beam and Rotarod assessments were performed, and pumps were replaced at times indicated. Mice were sacrificed after the final Rotarod evaluation at 36 weeks of age and extent of cerebellar pathology determined. By both balance beam (Figure 1B) and Rotarod (Figure 1C) assessments, two cohorts of *ATXN1[30Q]D776/Cck^-/-^* mice were equally compromised neurologically prior to initiation of treatment. Upon peripheral administration of either the Cck1R agonist A71623 or vehicle, *ATXN1[30Q]D776/Cck^-/-^* animals that received the A72613 agonist performed significantly better than *ATXN1[30Q]D776/Cck^-/-^* mice that received vehicle alone (Figure 1B and 1C). Purkinje cell pathology in the *ATXN1[30Q]D776/Cck^-/-^* animals receiving the Cck1R A72613 agonist, as assessed by extent of molecular layer atrophy and loss of Purkinje cells (Figure 1D), was significantly less than *ATXN1[30Q]D776/ Cck^-/-^* mice that received vehicle alone. Of note, while Cck has been shown to reduce food intake by activating Cck1R (reviewed in Miller and Desai, 2016), we found no difference in body weight between A71623 and vehicle treated *ATXN1[30Q]D776/Cck^-/-^* animals at 36 weeks-of-age (Figure S3).

**Figure 1.**
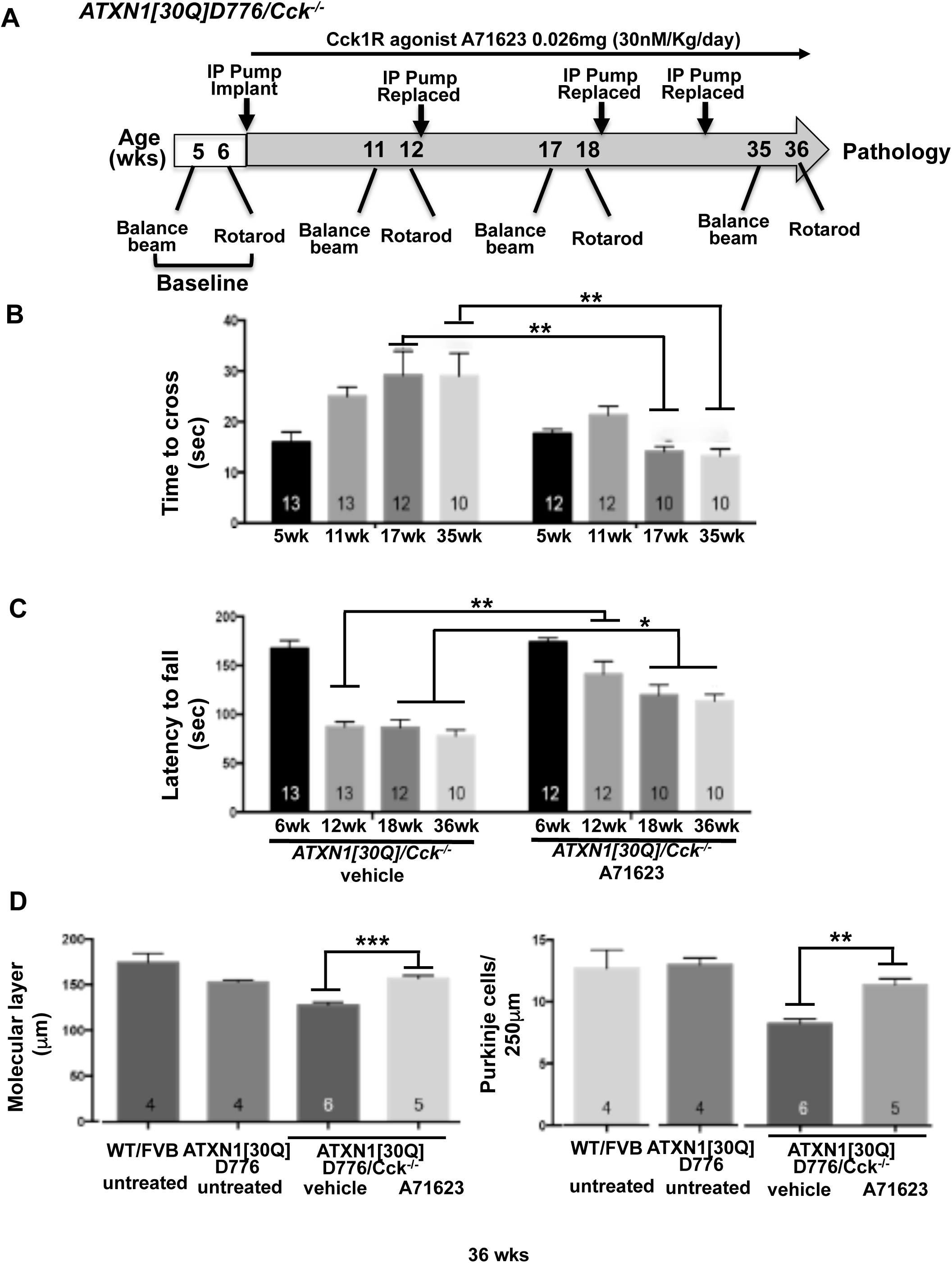
Cck1R agonist A71623 treatment dampens Purkinje neuron pathology in *ATXN1[30Q]D776;Cck^-/-^* mice. A) Scheme of assessment and treatment course. Mice were tested for motor performance using two tests. Then either 0.02mg/kg/Day A71623 or vehicle (20mM PBS) was administered until 36 weeks of age, at which time mice were sacrificed for pathology. Osmotic minipumps were replaced every 6 weeks. B) Time to cross the 10mm round balance beam. C) Latency to fall on the Rotarod. D) Molecular layer thickness in 36 week old untreated and treated mice. E) Number of Purkinje neurons per 250 um in cerebellar primary fissure in 36 week old untreated and treated mice. Two-Way ANOVA with Tukey post-hoc test, *p<0.05, **p<0.01, ***p<0.001. N’s (B-E) are indicated within each bar on the graphs.

### Overlap between *ATXN1[30Q]D776/Cck^-/-^* and *ATXN1[82Q]* Disease-Associated Gene Co-expression Modules

Previously we showed that a Weighted Gene Co-expression Network Analysis (WGCNA) of cerebellar RNA-seq data revealed a module reflecting Purkinje cell disease-correlated signature of gene expression alterations in *ATXN1[82Q]* animals (Ingram et al, 2016). To gain an understanding of the disease-correlated gene expression alterations in *ATXN1[30Q]/Cck^-/-^* mice, RNA-seq was performed on *ATXN1[30Q]/Cck^-/-^* and *Cck^-/-^* cerebella at 5, 12, and 28 weeks followed by WGCNA (Langfelder and Horvath, 2008). Two WGCNAs were performed; the first included the *ATXN1[30Q]/Cck^-/-^* RNA-seq data plus that obtained previously from WT, *ATXN1[30Q]D776, and ATXN1[82Q]* cerebella and a second one in which the *ATXN1[82Q]* RNA-seq data were not included in the WGCNA.

Figure 2A depicts the 36 WGCNA modules obtained on all RNA-seq data including that from *ATXN1[82Q]* cerebella. One module, the 374 gene Pink module, was found to significantly correlate with disease (p=7e-13) as determined by thickness of the cerebellar molecular layer. An assessment of the coexpression pattern of the Pink module using its eigengene showed that the eigengene in both *ATXN1[82Q]* and *ATXN1[30Q]D776/Cck^-/-^ and* not in *ATXN1[30Q]D776* decreased with age (Figure 2B) in a manner similar to the Magenta eigengene (Ingram et al., 2016). Consistent with the time course of molecular layer thinning in *ATXN1[82Q]* and *ATXN1[30Q]D776/Cck^-/-^* mice (Figure S1), the Pink eigengene decreased most pronouncedly between 5 and 12 weeks in *ATXN1[82Q]* and not until 28 weeks in *ATXN1[30Q]D776/Cck^-/-^* cerebella (Figure 2B). Interestingly, the PINK WGCNA module obtained using *ATXN1[30Q]D776/Cck^-/-^* RNA-seq data overlapped extensively and to a highly significant level with the previous disease-associated Magenta WGCNA module (Ingram et al, 2016) (Figure 2C). To rule out that this overlap was driven by *ATXN1[82Q]* RNA-seq data, a second WGCNA was performed excluding the *ATXN1[82Q]* data. Again a Pink module was found to significantly correlate with molecular layer thickness that also overlapped highly significantly (p=1.4e-103) with the previous Magenta WGCNA module (Figure 2C). Thus, we conclude that the RNA-seq/WGCNA of *ATXN1[82Q]* and *ATXN1[30Q]D776/Cck^-/-^* supports the conclusion that the Purkinje cell disease in these two *ATXN1* transgenic lines shares an underlying molecular basis.

**Figure 2.**
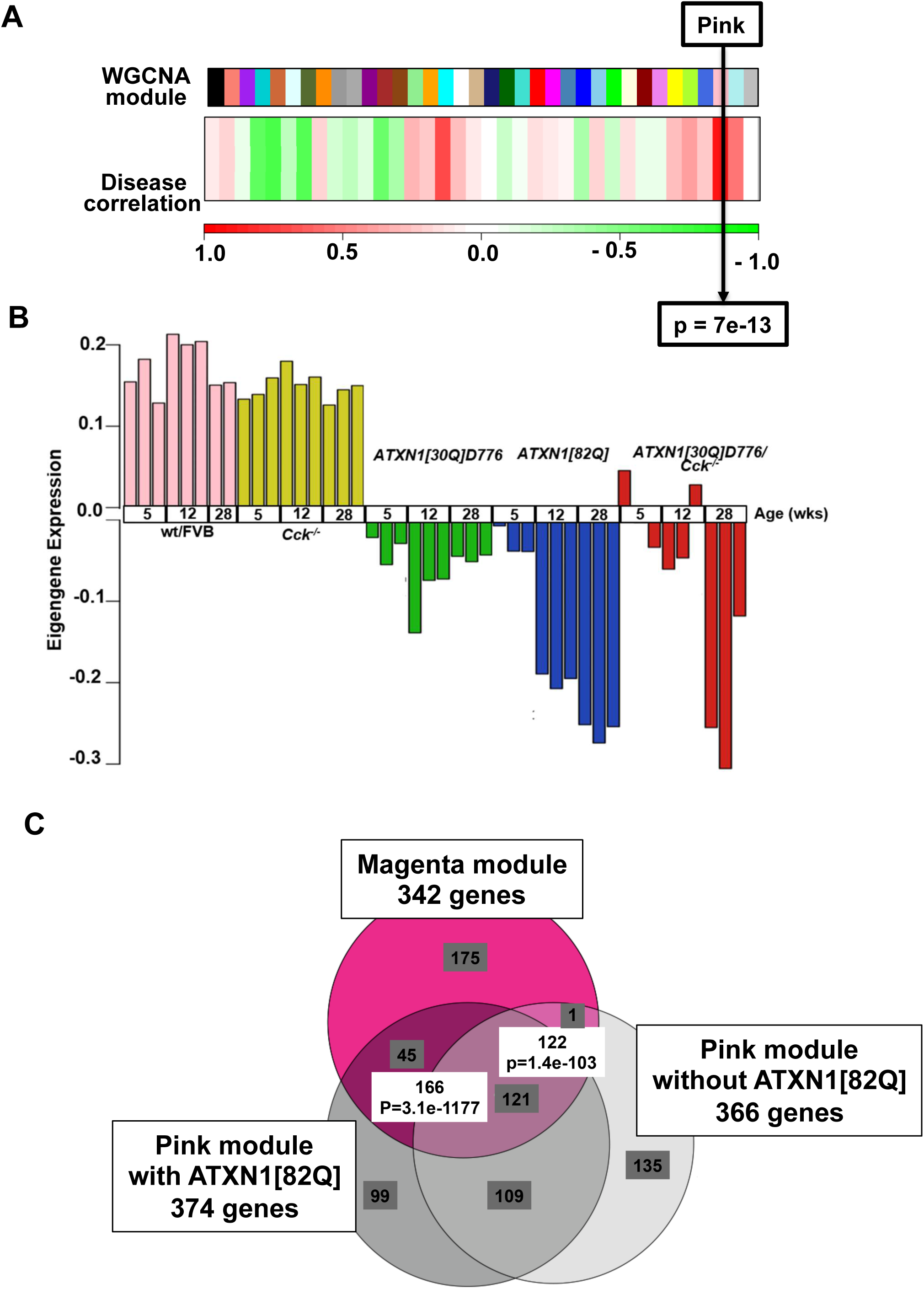
Cerebellar WGCNA for *ATXN1[30Q]D776;Cck^-/-^* Mice. A) *ATXN1[30Q]D776;Cck^-/-^* cerebellar WGCNA including all samples reveals one gene module (Pink) that correlates significantly with molecular layer thickness across all ages. Fischer’s Exact test followed by Benjamin Hodgeberg adjustment to control false discovery rate. B) Pink module eigengene expression reveals similar genes expression changes in ATXN1[82Q] and *ATXN1[30Q]D776;Cck^-/-^* samples across time. C) Venn diagram depicting overlap in DEGs between *the ATXN1[30Q]D776;Cck^-/^-* WGCNA Pink and the *ATXN1[82Q*] WGCNA Magenta modules

### Cck1R Agonist A71623 Alleviates Disease in *ATXN1[82Q]* Mice

With the significant similarity in age-related gene co-expression changes in *ATXN1[82Q]* and *ATXN1[30Q]D776/Cck^-/-^* cerebella along with a decrease in *Cck* expression in Purkinje cells of *ATXN1[82Q]* mice (Ingram et al., 2016), we reasoned that the Cck1R agonist A71623 would likely dampen disease in *ATXN1[82Q]* mice. Thus, a treatment scheme based on that used to show the ability of the Cck1R agonist A71623 (dosed at 0.026mg/Kg/day) to impact Purkinje cell disease progression in *ATXN1[30Q]D776/Cck^-/-^* mice was applied to *ATXN1[82Q]* mice (Figure 3A). Due to the more progressive nature of disease in *ATXN1[82Q]* animals, the strategy utilized included two modifications from that used with *ATXN1[30Q]D776/Cck^-/-^* mice. First, agonist treatment was initiated earlier at 4 weeks of age. Therefore, until animals reached a size such that the IP pumps could be implanted, they were given IP injections of A71623 for two weeks. The second modification was that the initial trial was run for 12 weeks, to the age when *ATXN1[82Q]* mice show the most dramatic decrease in the thickness of the molecular layer (Figure S1). Following a final motor assessment at 11 and 12 weeks of age, A71623 and vehicle treated animals were sacrificed and analyzed using molecular and morphological approaches.

**Figure 3.**
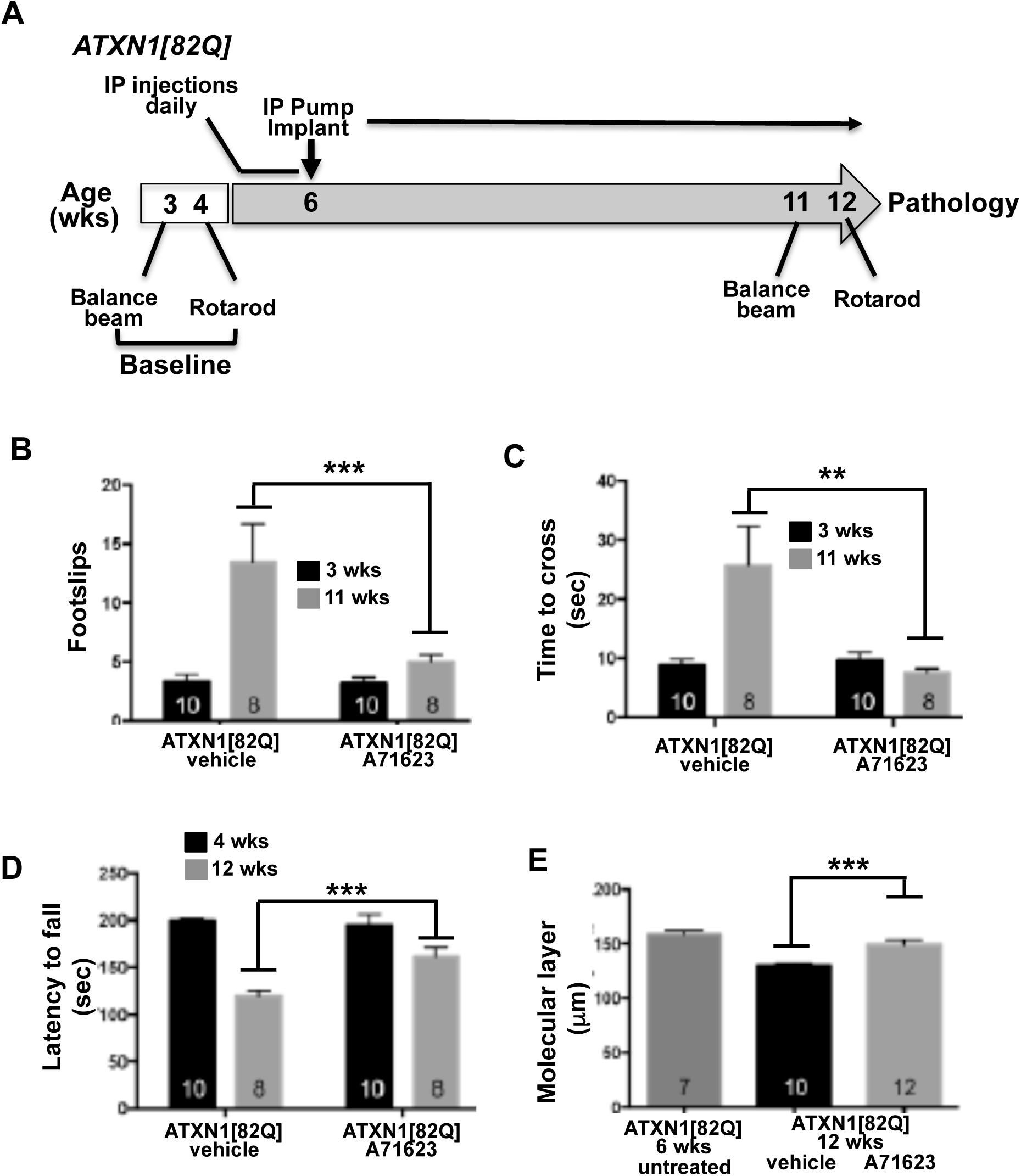
Cck1R agonist A71623 treatment dampens Purkinje neuron pathology *ATXN1[82Q]* mice. A) Scheme of the assessment and treatment course. Mice were tested for motor performance using two tests. Then either 0.02mg/kg/Day A71623 or vehicle (20mM PBS) was given until 12 weeks of age, at which time mice were sacrificed for pathology. B) Number of footslips on the 10mm round balance beam. C) Time to cross the 10mm round balance beam. D) Latency to fall on the Rotarod. E) Molecular thickness of 6-week-old untreated mice compared to 12 week old treated mice. Two-Way ANOVA with Tukey post-hoc test, *p<0.05, **p<0.01, ***p<0.001.

By beam walk assessments (Figures 3B and 3C) assessing footslips and time to cross a 10mM round beam, respectively, the cohorts that were subsequently treated with A71623 or vehicle were equally impaired at 3 weeks of age. However, at 11 weeks of age, following 7 weeks of treatment, *ATXN1[82Q]* mice treated with the Cck1R agonist A71623 performed significantly better on the beam than those administered vehicle alone. Likewise, Rotarod evaluation indicated that both cohorts performed equally well at 4 weeks of age. Yet, by 12 weeks of age, performance on the Rotarod of vehicle treated *ATXN1[82Q]* mice deteriorated more significantly than *ATXN1[82Q]* animals that received the agonist A71623 (Figure 3D). Similarly, molecular layer thinning at 12 weeks of age was significantly worse in vehicle treated compared to A71623 treated *ATXN1[82Q]* mice (Figure 3E). Importantly, the mechanism by which A71623 treatment alleviated disease in the ATXN1[82Q] mice did not involve a reduction in levels of the ATXN1[82Q] protein (Figure S4).

### Cck1R Agonist A71623 Improves Motor Performance of *ATXN2[127Q]* Mice

To assess whether the Cck1R agonist A71623 has a protective effect on another SCA mouse model, we utilized a mouse model of SCA2 expressing *ATXN2[127Q]* under the Purkinje cell-specific regulatory element from the *Pcp2* gene (Hansen et al., 2013). The treatment scheme and dosage for the Cck1R agonist A71623 was identical to that used for *Pcp2-ATXN1[82Q]* mice (Figure 4A). At 11 weeks of age, *Pcp2-ATXN2[127Q]* mice had a significant deficit in performance on the balance beam by both number of footslips and time to cross (Figure 4B and 4C). Importantly, at 11 weeks of age following 7 weeks of treatment *Pcp2-ATXN2[127Q]* mice treated with the Cck1R agonist A71623 performed significantly better on the balance beam than those administered vehicle alone (Figure 4B). On the Rotarod at 12 weeks of age the *Pcp2-ATXN2[127Q]* mice showed no signs of a progressive deficit in performance compared to their performance at 4 weeks of age relative to age-matched WT/FVB mice (data not shown).

**Figure 4.**
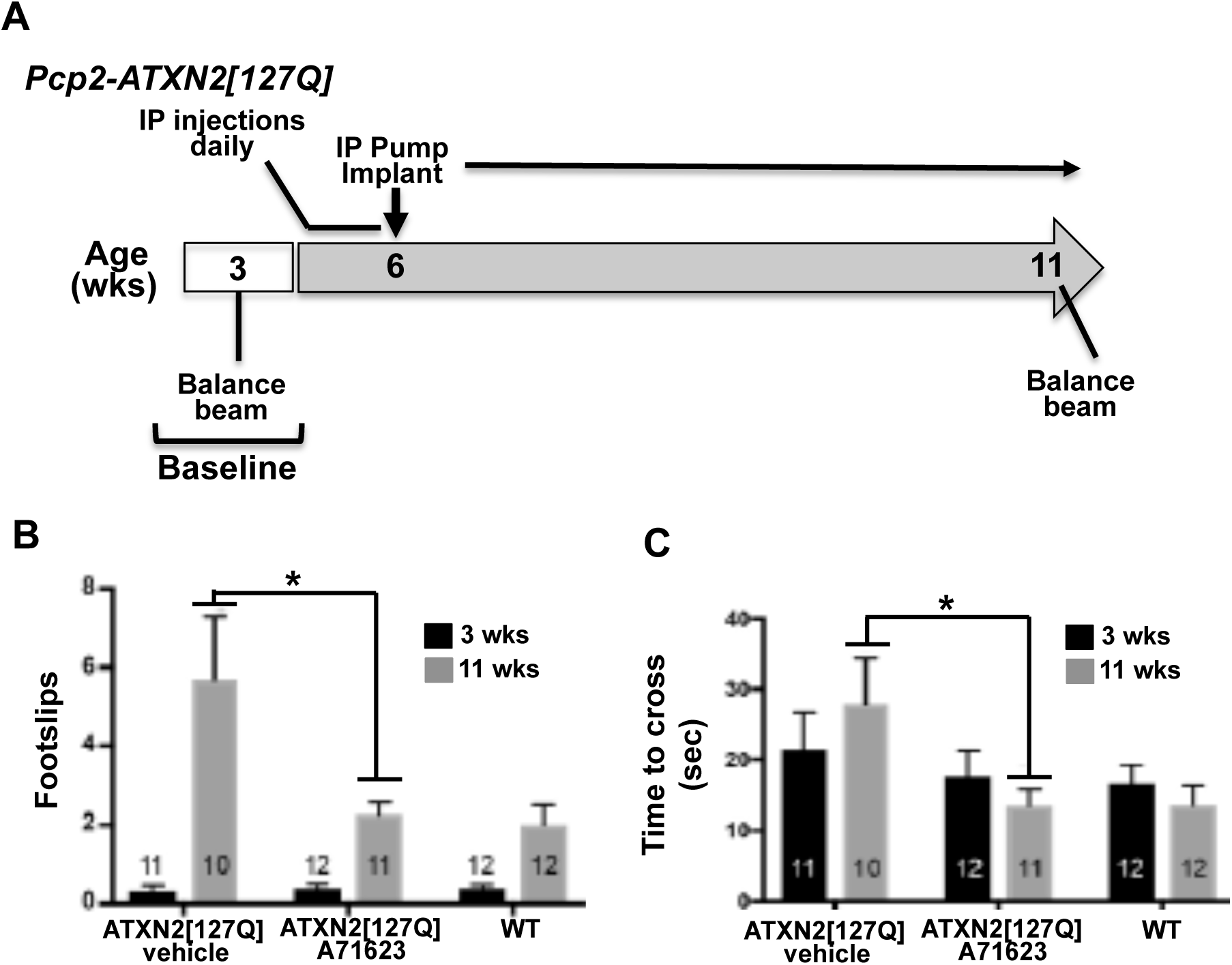
Cck1R agonist A71623 treatment improves motor performance in *ATXN2[127Q]* mice. A) Schematic depiction of the assessment and treatment course. Mice were tested for motor performance, baseline at 3 weeks of e and at 11 weeks of age using the 10mm round balance beam. At 4 weeks of age mice were given daily IP injections of 0.02mg/kg/day A71623 or vehicle (20mM PBS). At 6 weeks of age osmotic pumps were implanted intraperitoneally and the Cck1R agonist or vehicle were administered. B) Number of footslips on the 10mm round balance beam. C) Time to cross the 10mm round balance beam. Two-Way ANOVA with Tukey post-hoc test, *p<0.05.

### Proper mTORC1 Signaling is Restored by Cck1R Agonist A71623 in *ATXN1[82Q]* and *ATXN1[127Q]* Cerebella

Ruegseggar et al., demonstrated that reduced mTORC1-signaling contributes to Purkinje cell disease in the cerebellum of the *Atxn1^154Q/2Q^* knockin mouse model of SCA1 (Ruegseggar et al., 2016). Importantly, mTORC1 is a downstream signaling pathway activated via Cck1R stimulation (reviewed in Cawston & Miller, 2010). Thus, we assessed the effect of the Cck1R agonist A71623 on mTORC1 signaling status in *Pcp2-ATXN1[82Q]* and *Pcp2-ATXN2[127Q]* cerebella. Figure 5A shows that mTORC1 signaling was significantly decreased in 11-week-old *ATXN1[82Q]* cerebella as evaluated by measuring the phosphorylation level of the ribosomal protein S6. Next, we examined whether administration of the Cck1R agonist A71623 restored mTORC1 signaling in *ATXN1[82Q]* cerebella. Six-week-old *ATXN1[82Q]* mice were given an IP injection of A71623 (0.264mg/kg). Twenty-four hours later, cerebella were harvested and mTORC1 signaling was found to be restored to a WT level as assessed by phosphorylation of the ribosomal protein S6 (Figure 5B). Consistent with the ability of A71623 to decrease pathology in ATXN1[82Q] mice (Figure 3) a small but significant increase in the amount of calbindin protein was detected cerebellar extracts from A71623 treated mice (Figure 5B). To validate that the action of A71623 on mTORC1-signaling were via the Cck1R, *ATXN1[82Q]* mice were crossed to Cck1R^-/-^ mice (Kopin et al., 1999). Figures 5C and 5D show that in eight-week-old *ATXN1[82Q]* mice lacking Cck1R, A71623 was no longer able to restore cerebellar mTORC1 signaling.

**Figure 5.**
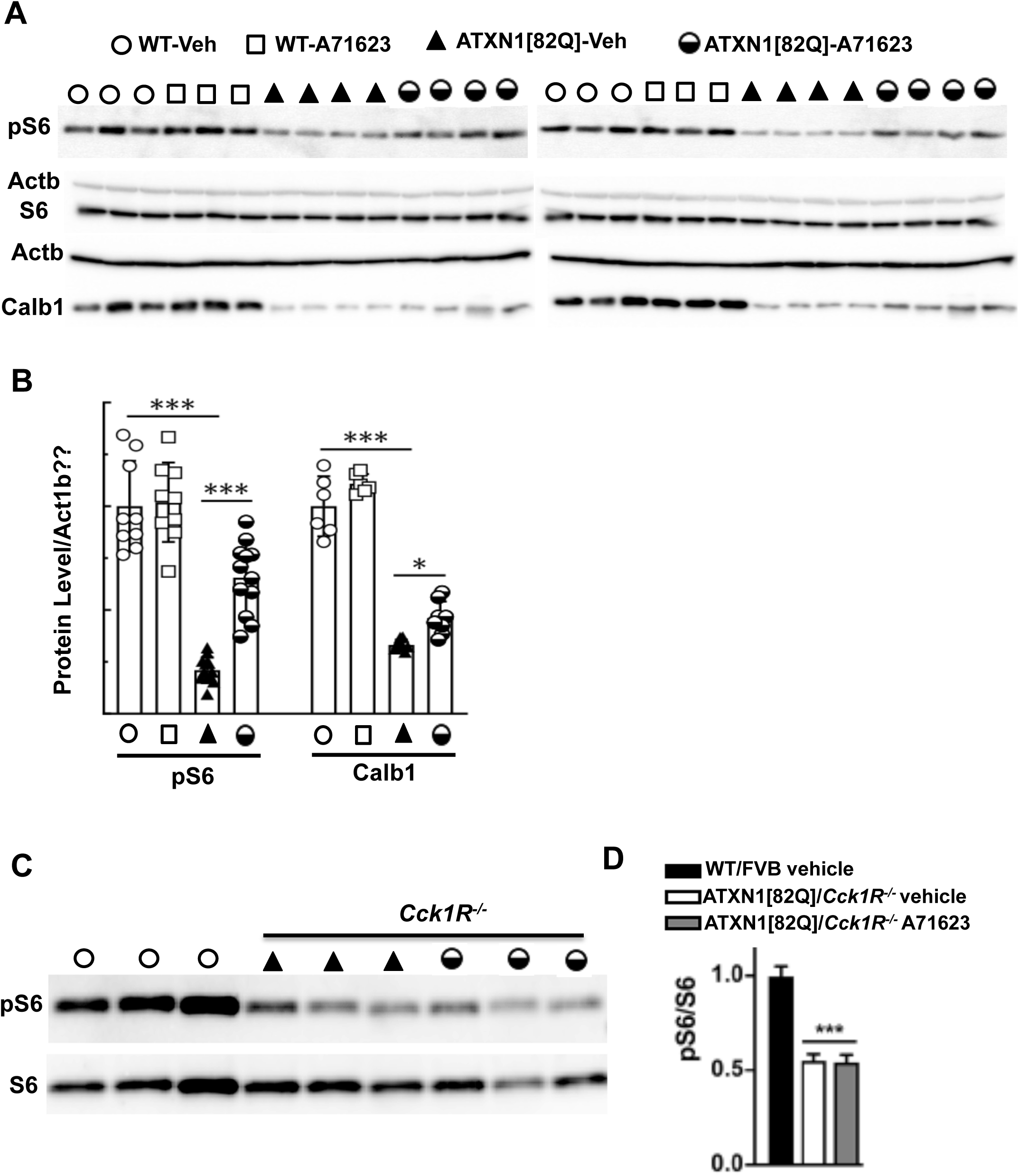
mTORC1 signaling activity and cerebellar marker Calb1 are reduced in ATXN1[82Q] mice and restored by Cck1R activation. A) Phosphorylation of the ribosomal protein S6 (pS6) is significantly reduced in the cerebellum *of ATXN1[82Q]* mice (11 wks of age) compared to WT/FVB control mice. Administration of the Cck1R-selective agonist A71623 to *ATXN1[82Q]* mice for 24 hr restores cerebellar phosphorylation of S6 and Calb1. Each lane represents extract from an individual mouse. Actb is used as a loading control and the blots are from replicate experiments. B) Quantification of P-S6 and Calb1 expression levels in cerebella of *ATXN1[82Q] mice* . C) Absence of the Cck1R in *ATXN1[82Q]* mice prevents A71623 restoration of cerebellar S6 phosphorylation. One-way ANOVA followed by Bonferroni’s multiple comparisons test. Data are mean ± SD, ns = *P* > 0.05, \**P* < 0.05, \*\**P* < 0.01, *\*\*\*P* < 0.001.

Figure 6 presents shows that the Cck1R agonist A71623 also normalized cerebellar mTORC1 signaling in *ATXN2[127Q]* mice. In contrast to *ATXN1[82Q]* where mTORC1 signaling is decreased, *ATXN2[127Q]* mice show hyperactivation of mTOR signaling resulting in autophagy aberration (Paul et al., 2021). However, both mouse models share similar cerebellar marker gene dysregulations including Calb1. To investigate if Cck1R agonist (A71623) has effect on mTORC1 signaling, we performed intraperitoneally (IP) bolus injection of A71623 (0.0264, 0.264 and 1.0 mg/kg) in *ATXN2[127Q]* mice at 13 weeks of age for 24 hr. On western blot analyses, *ATXN2[127Q]* mice showed increased phospho-S6 ribosomal protein levels as a function of mTORC1 activation (**Figure 6A, B**). CcK1R agonist treatment successfully reduced P-S6 levels to wildtype control levels (**Figure 6A, B**). Complementary to reduced pS6 levels, we also observed significant restoration of cerebellar gene, Calb1 in *ATXN2[127Q]* mice by Cck1R agonist treatment (**Figure 6A, B**). Quantification of pS6 and Calb1 protein levels of three independent blots are shown in **Figure 6C**.

**Figure 6.**
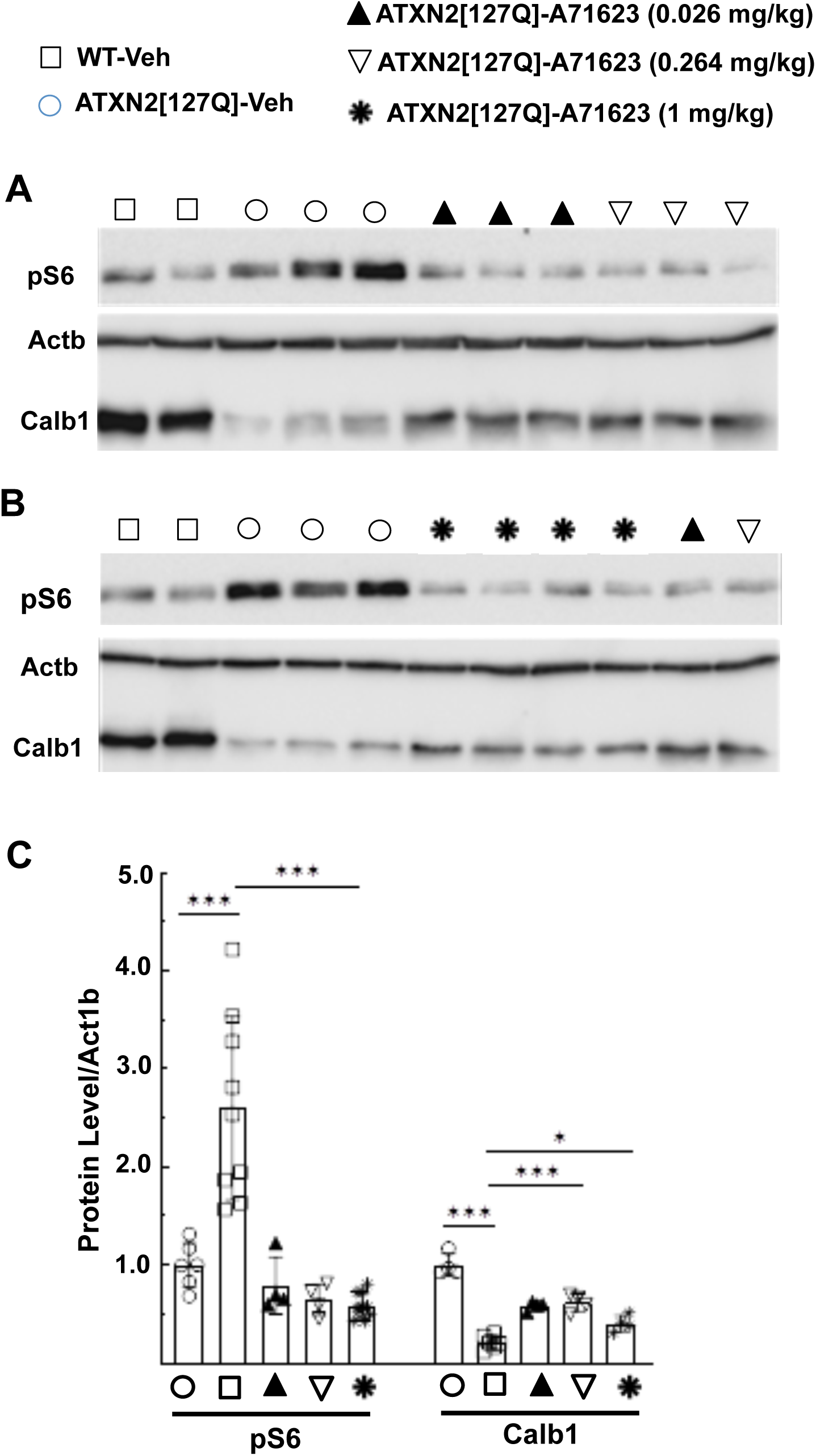
Cck1R activation normalizes mTORC1 signaling activity and restores cerebellar marker Calb1 in ATXN2[Q127] mice. A) Phosphorylation of the ribosomal protein S6 (pS6) is increased and cerebellar marker Calb1 is decreased in the cerebellum *of ATXN2[127]* mice (13 wks of age) compared to wildtype mice. Administration of the Cck1R-selective agonist A71623 to *ATXN2[127Q]* mice for 24 hr restores cerebellar P-S6 and Calb1. Each lane represents extract from an individual mouse. Actb is used as a loading control and blots are from replicate experiments. B) Quantification of P-S6 and Calb1 are shown. One-way ANOVA followed by Bonferroni’s multiple comparisons test. Data are mean ± SD, ns = *P* > 0.05, \**P* < 0.05, \*\**P* < 0.01, *\*\*\*P* < 0.001.

## DISCUSSION

In many SCAs, cerebellar Purkinje neurons manifest pathology and dysfunction (Robinson et al., 2020). Previous RNA-seq analyses of mouse models of SCA1 revealed an enhanced expression of *Cck* in cerebellar RNA from *ATXN1[30Q]D776* mice that manifest severe ataxia in absence of progressive Purkinje cell pathology (Ingram et al. 2016). Development of ataxia with progressive Purkinje cell pathology upon crossing *ATXN1[30Q]D776* mice with *Cck^-/-^* and *Cck1R^-/-^* mice suggested that the elevated Cck in *ATXN1[30Q]D776* mice and its activation of Cck1R brought about the lack of Purkinje cell pathology in these mice. Here we show that a Cck1R-selective agonist A71623 was able to mitigate the ataxia and progressive Purkinje cell pathology in *ATXN1[30Q]D776/Cck^-/-^*, *ATXN1[82Q]*, and *ATXN2[127Q]* mice, providing direct evidence that activation of Cck1Rs is neuroprotective in SCA cerebellar disease.

CCK was first identified as a gastrointestinal tract hormone regulating GI motility, pancreatic enzyme secretion, gastric emptying, and gastric acid secretion. Subsequently CCK was found to be a widely expressed and abundant neuropeptide in the CNS where it is reported to regulate output of neuronal circuits, notably satiation (Lee and Slotesz, 2011). Co-localization and interaction of CCK with other key neurotransmitters in CNS areas suggests that CCK has an extensive role regulating neuronal activity. CCK interacts with two G-protein coupled receptors to mediate its biological actions; CCK1R and CCK2R (Dufresne et al., 2006). In the cerebellum, a prominent site of *Cck* and *Cck1R* expression are Purkinje neurons (Figure S5).

Typically, the mTORC1 signaling pathway is viewed as being inhibited by varous stresses (reviewed in Su and Dai, 2017). Based on the results presented in this study along with our previous work (Ingram et al., 2016), in the case of ATXN1[82Q], we propose a model whereby a CCK/CCK1R/mTORc1 pathway has a role in Purkinje cells adjusting to stress and that this pathway is negatively impacted by ATXN1 with an expanded polyQ tract (Figure 7). In ATXN1[82Q] Purkinje cells, stress promotes the cleavage of CCK to the octapeptide CCK-8, the natural ligand with the highest affinity for the Cck1R (Dufresne et al., 2006) that upon secretion binds to and activates CCK1R on Purkinje cells. Expression of ATXN1[82Q] enhances Purkinje cell stress. The fact that ATXN1 with an expanded PolyQ also reduces CCK expression dampens the ability of Purkinje cells to respond to stress thus promoting ATXN1 pathogenesis. Activation of Purkinje cell CCK1R stimulates mTORC1, which we speculate is a critical component by which CCK1R activation dampens the pathogenic effects of ATXN1 with an expanded PolyQ. Inactivation of mTORC1 induces a progressive loss of Purkinje cells by apoptosis (Angliker et al., 2015). mTORC1 signaling, in addition to responding to many stresses, is impaired in ATXN1^154Q^ knockin mice and absence of mTORC1 in Purkinje cells of ATXN1^154Q^ mice worsens disease (Ruegsegger et al., 2016).

**Figure 7.**
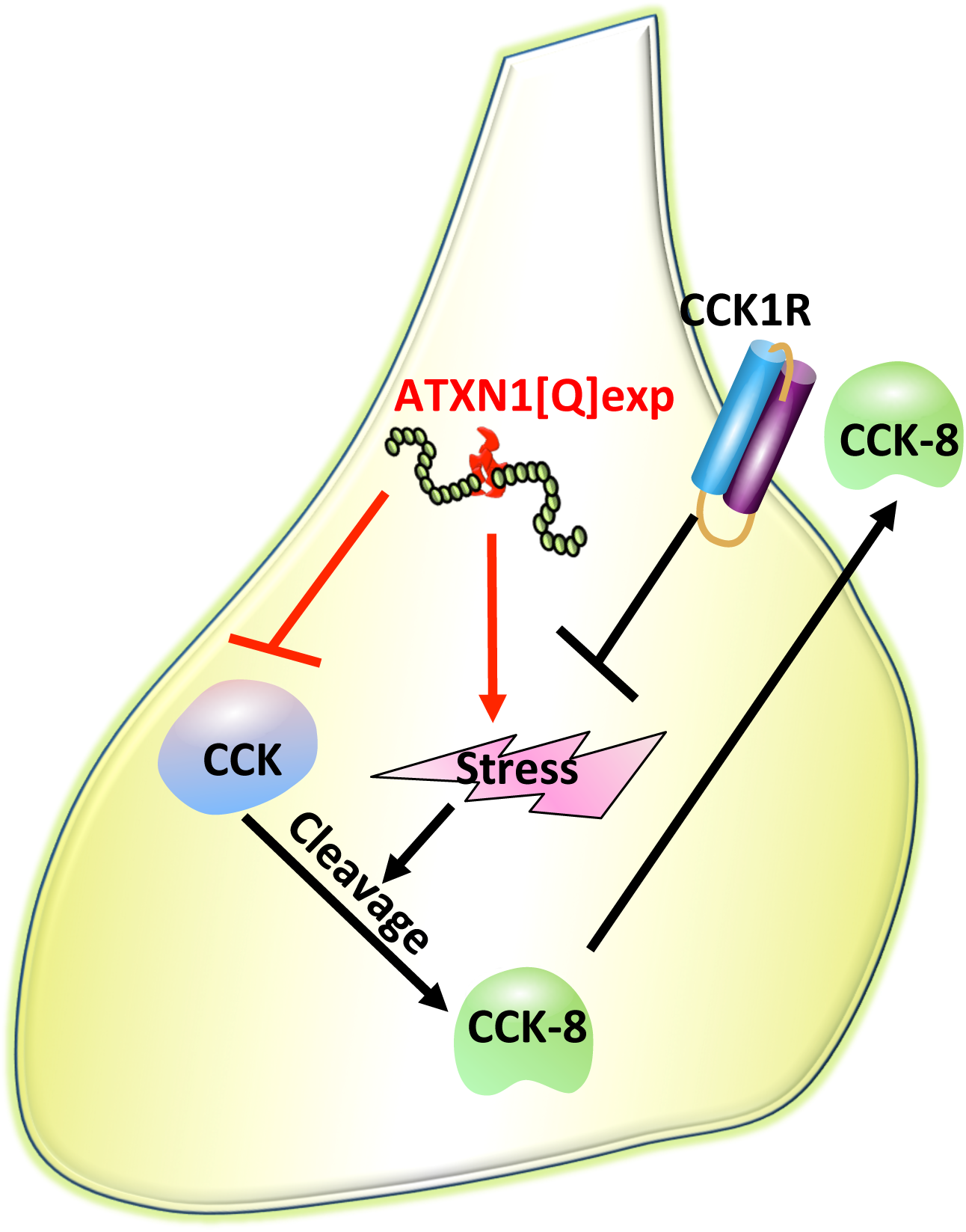
A Cck1R activated mTORC1 pathway that responds to stress in Purkinje cells and is diminished by ATXN1 with an expanded polyQ. In response to stress Purkinje cells increase the cleavage of Cck to Cck-8 that is secreted and binds to and activates Cck1Rs located on the cell membrane of Purkinje cells. Activation of Cck1Rs promotes mTORC1 signaling thus promoting Purkinje cell homeostasis. As indicated in red, ATXN1 with an expanded polyQ interferes with thisCck1R-mTORC1 stress response pathway by both increasing the amount of stress in Purkinje cells as well as decreasing the expression of Cck, resulting in a reduction in Purkinje cell homeostasis.

In contrast to ATXN1[82Q] expressing Purkinje cells where mTORC1 signaling is decreased, in *ATXN2[127Q]* Purkinje cells have enhanced mTORC1 signaling. Previously, Angliker et al. 2015 noted a striking similarity in the effect between Purkinje cells with decreased mTORC1-signaling compared to those in which mTORC1-signaling was enhanced. In mice where mTORC1-signaling was decreased due to a loss of *mTORC1* or in Purkinje cells in which mTORC1-signaling was enhanced due to the absence of the mTORC1 inhibitor *TSC1*, the function as well as survival of Purkinje cells were adversely affected. The data presented here on mTORC1-signaling levels in cerebella of *ATNX1[82Q]* and *ATXN2[127Q]* transgenic mice indicate that SCAs might also be categorized according to mTORC1-signaling activity with SCA1 having reduced Purkinje cell mTORC1-signaling and SCA2 being a SCA with enhanced Purkinje cell mTORC1-signlaing. Intriguingly, activation of the Cck1R in both instances restored mTORC1-signlaing to a normal level. Understanding the mechanism and cellular pathways by which Cck1R activation in ATNX1[82Q] and *ATXN2[127Q]* transgenic Purkinje cells restores mTORc1-signlaing to a normal level is an area of considerable importance regarding the potential of Cck1R activation as a therapeutic target in the SCAs.

## ACKNOWLEDGEMENTS

This study was supported by NIH/NINDS grant RO1NS 045667 (H.T.O.), and NIH/NINDS grants R37NS033123, R21NSNS103009 and UO1NS103883 (SMP). The authors thank the Biomedical Genomics Center, Mouse Phenotyping Core at the University of Minnesota and Orion Rainwater for propagating and maintaining mouse colony.

## AUTHOR CONTRIBUTIONS

E.A.L.W.,Z.C. and S.P. contributed equally to this work. E.A.L.W., Z.C., S.P., K.P.F, S.M.P. and H.T.O. contributed to the conception and design of the study. E.A.L.W., Z.C., P.Y., J.F. T.T. and M.B. contributed to tissue selection, collection and analyses. E.A.L.W. and M.I. performed RNA extractions for RNA-seq. M.I., and C.H., performed RNA-seq expression and WGCNA bioinformatics analyses. C.H. supervised bioinformatics analyses. M.I., E.A.L.W., Z.C., S.P., P.Y., C.H., S.M.P. and H.T.O. interpreted the data and prepared the manuscript.

## EXPERIMENTAL PROCEDURES

### Mice

The Institutional Animal Care and Use Committee approved all animal use protocols. *ATXN1[82Q]* (Burright et al., 1995), *ATXN1[30Q]-D776, ATXN1[82Q]-D776* (Duvick et al., 2010), *ATXN2[127Q]* (Hansen et al., 2013), and WT/FVB/NJ mice were housed and managed by Research Animal Resources under SPF conditions in an AAALAC-approved facility. In all experiments involving the use of mice, equal numbers of male and females were used.

### RNA isolation and sequencing

Total RNA was isolated from dissected cerebella using TRIzol Reagent (Life Technologies, Carlsbad, California) following the manufacturer’s protocols. Cerebella were homogenized using an RNase-Free Disposable Pellet Pestles in a motorized chuck. For RNA-sequencing, RNA was further purified to remove any organic carryover using the RNeasy Mini Kit (Qiagen, Venlo, Netherlands) following the manufacturer’s RNA Cleanup protocol.

Cerebellar RNA from three biological replicates for each genotype was isolated. Purified RNA was sent to the University of Minnesota Genomics Center for quality control, including quantification using fluorimetry (RiboGreen assay, Life Technologies) and RNA integrity assessed with capillary electrophoresis (Agilent BioAnalyzer 2100, Agilent Technologies, Inc.) generating an RNA integrity number (RIN). All submitted samples had greater than 1ug total mass and RINs 7.9 or greater. Library creation was completed using oligo-dT purification of polyadenylated RNA, which was reverse transcribed to create cDNA. cDNA was fragmented, blunt-ended, and ligated to barcoded adaptors. Library was size selected to 320bp +/-5% to produce average inserts of approximately 200bp, and size distribution validated using capillary electrophoresis and quantified using fluorimetry (PicoGreen, Life Technologies) and Q-PCR. Libraries were then normalized, pooled and sequenced. 12 and 28 week *ATXN1[82Q]*, *ATXN1[30Q]-D776*, and WT/FVB samples were sequenced on an Illumina GAIIX using a 76nt paired-end read strategy, while 5 week samples from these genotypes were sequenced on an Illumina HiSeq 2000 using a 100nt paired-end read strategy. Data were stored and maintained on University of Minnesota Supercomputing Institute (MSI) Servers.

### Gene expression analyses

Gene expression analyses were performed with the Tuxedo pipeline (Kim et al., 2013b; Trapnell et al., 2010). Initial read quality was assessed using FastQC (Andrews - Babraham Bioinformatics, FastO|QC A quality control tool for high throughput sequence data) and reads were trimmed to remove low quality 3’ ends and adapter contamination using Trimmomatic (Bolger et al., 2014).

Reads were aligned to the mouse reference genome (mm10) with Tophat by using mostly default parameters with two exceptions: mate inner distance and standard deviation were adjusted to the data and a gene annotation model only looking for supplied junctions (mm10 gtf file from iGenomes). Differential gene expression was determined with Cuffdiff using default parameters (Trapnell et al, 2012). Genes/introns with a q≤0.05 were considered significant. Genome tracks were visualized with Integrated Genomics Viewer (Broad Institute). Results were graphed with CummeRbund (http://compbio.mit.edu/cummeRbund/; Goff et al., 2013).

### WGCNA

FPKM abundance estimates for all 27 samples were produced by CuffNorm (Trapnell et al. 2012) and were log_2_ transformed (log_2_(FPKM+1)) for WGCNA analysis (Langfelder and Horvath, 2008). The WGCNA R package (v. 1.41) was used to construct an unsigned gene coexpression network with a soft threshold power [beta] of 10. Nineteen modules were detected, including two that were significantly associated with ataxia (WT vs *ATXN1[82Q]* and *ATXN1[30Q]D776* mice, t-test, Bonferroni corrected p-value < 1e-5).

### Agonist treatments

#### Osmotic pumps

The Cck1 receptor (Cck1R) agonist A71623 (Tocris Biosciences) was suspended in 20mM PBS according to the manufacturer’s directions. For the experiment with *ATXN1[30Q]D776;Cck^-/-^* mice (Figure 14B), osmotic minipumps (Alzet, 1004) containing either A71623 (0.02mg/kg/day) or Vehicle (20mM PBS) were implanted intraperitoneally (i.p.) in 6 week old mice. Briefly, 5 week old mice were deeply anesthetized by intramuscular injection of a ketamine/xylazine cocktail (100mg/kg ketamine and 10mg/kg xylazine). Fur on i.p. implantation site (1cm below ribcage) was shaved. Incision sites were scrubbed with povidone-iodine (Betadine) scrub. A 1cm-long incision was made under the ribcage. The peritoneal wall was gently incised beneath the cutaneous incision and the pump cannula was placed into the peritoneal cavity. The musculoperitoneal layer was closed with 4.0 absorbable suture and the skin wound was closed using surgical staples (Alzet). Ten days after surgery, the staples were removed and wounds examined for healing. For the duration of the experiment, pumps were removed and replaced every 7 weeks. Behavioral data was collected at the time points indicated in Figure 4B. Because of the size of the osmotic minipumps, the mice have to be ∼20g or larger for safe implantation into the i.p. space. In ATXN1[82Q] mice, single bolus injections of A71623 (0.02mg/kg) or Vehicle were administered daily beginning at week 5 and continuing until the mice were ∼20g (for approximately 7 days, or until the mice were 6 weeks old). At this time the pumps were implanted for the remainder of the experimental timeline.

#### I.P. injections

*ATXN2[127Q]* mice (13 weeks of age) were given intraperitoneally (IP) bolus injection of Cck1R agonist (A71623: 0.0264, 0.264 and 1.0 mg/kg) or vehicle (20 mM PBS) for 24 hr. Following A71623 treatment, cerebella were harvested and stored at −80°C until used.

### Western blot

Mouse cerebellar extracts were prepared by homogenization of tissues in extraction buffer [25 mM Tris-HCl pH 7.6, 250 mM NaCl, 0.5% Nonidet P-40, 2 mM EDTA, 2 mM MgCl2, 0.5 M urea and protease inhibitors (Sigma-Aldrich, P-8340)] followed by centrifugation at 4°C for 20 min at 14,000 RPM (Paul et al., 2021). Only supernatants were used for western blotting. Protein extracts were resolved by SDS-PAGE and transferred to Hybond P membranes (Amersham Bioscience, USA), and processed for western blotting. Immobilon Western Chemiluminescent HRP Substrate (EMD Millipore, Cat# WBKLSO500) was used to visualize the signals, which were detected on the ChemiDoc MP imager (Bio-Rad) and the band intensities were quantified by ImageJ software analyses after inversion of the images. Relative protein abundances were expressed as ratios to Actb. Antibodies used for western blotting and their dilutions were as follows: Phospho-S6 Ribosomal Protein (Ser235/236) (D57.2.2E) XP^®^ Rabbit mAb [(1:4000), Cell Signaling, Cat#4858], S6 Ribosomal Protein (5G10) Rabbit mAb [(1:4000), Cell Signaling, Cat# 2217], monoclonal anti-Calbindin-D-28K antibody [(1:5,000), Sigma-Aldrich, C9848], monoclonal anti-β-Actin−peroxidase (clone AC-15) [(1:30,000), Sigma-Aldrich, A3854]. Secondary antibodies: Peroxidase-conjugated AffiniPure goat anti-rabbit IgG (H + L) [(1:5000), Jackson ImmunoResearch Laboratories, Cat# 111-035-144] and goat anti-mouse IgG (Fab specific)–Peroxidase [(1:5000), Sigma-Aldrich/Millipore, Cat# A2304-1ML].

### Behavioral analyses

<u>Rotarod</u>. Mice were tested on the rotarod apparatus (Ugo Basile, Comerio, Italy) using an accelerating protocol; 5 to 50 rpm, 5 min ramp duration, 5 min max trial length. The test consisted of a total of 4 trials per day for 4 consecutive days. Mice were habituated to the testing room 15 minutes prior to the start of testing on each day. Animals were segregated by sex during testing and run in consistent groups (up to 5 at a time). To ensure enough recovery time between trials, animals were given 10-15 minutes between the end of a trial and the following trial, which included the time to test the other groups in the trial. Trials ended whenever an animal failed to stay on the rotarod or if they made 2 consecutive rotations clinging to the rod and not ambulating. Animals were returned to their home cages after trial completion. The apparatus was cleaned between each animal group within a trial with a 70% ethanol.

#### Beam walk

The beam walk protocol consisted of 3 consecutive training days followed by one test day. The beam walk apparatus was built in-house by the Mouse Behavior Core using wood and plastic components, consisting of beams (of varying sizes and shapes) that extend to a goal box. The apparatus is elevated ∼50 cm off a table surface. Testing was performed in a dark room with only a single light directed at the start position of the beam to encourage crossing to the goal box. Males and females were segregated during testing. Animals were run at approximately the same time every day and habituated to the behavior room for about 15 minutes prior to the start of training. At the start of the first trial of each day, an animal was placed into the goal box for ∼15-30 seconds to become familiar before beginning the first trial. After familiarization with the goal box, the mouse was brought to the start end of the beam and placed with the nose right behind the ‘start’ line. Beam walk training (using the 15 mm square beam) consisted of 4 trials per day, with a maximum of 60 seconds allowed for an animal to cross. Animals were tested in smaller groups of 4-7 to allow for ∼5 min intertrial interval. Hindpaw foot slips and time (sec) to cross the beam were carefully recorded by an investigator blind to treatment and genotype. A foot slip was defined as a hind leg and paw coming completely off the beam in a fast downward sweeping motion. Animals that successfully crossed the beam were left in the goal box for ∼15 seconds while the beam was wiped down with a 70% ethanol solution. The animal was then returned to their home cage until the next trial. The goal platform was cleaned with the ethanol solution after each trial and between animals on the first trial of each day.

---

**Figure S1.**
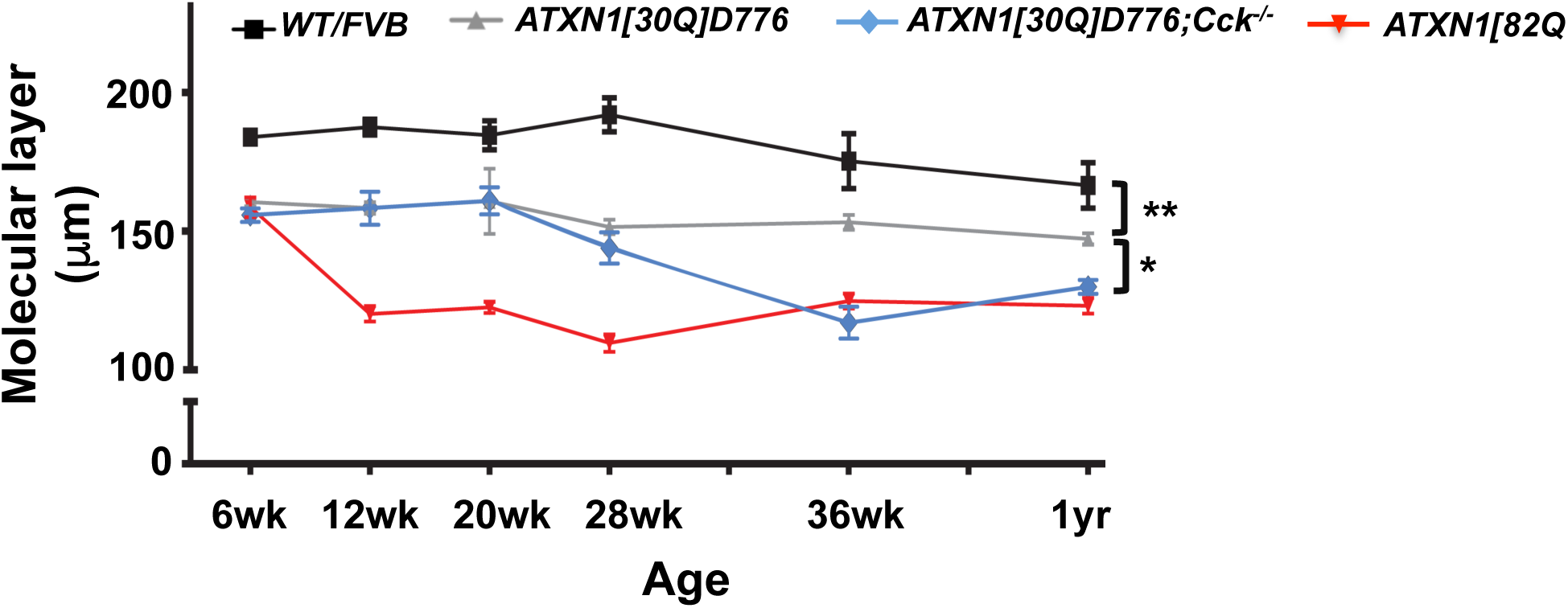
Purkinje cell Atrophy in ATXN1 Mice with Age. Purkinje Cell atrophy was assessed by measuring thickness of the cerebellar molecular layer. n>6 per group Two-way ANOVA, tukey post-hoc test. *p<0.05 **p<0.01.

**Figure S2.**
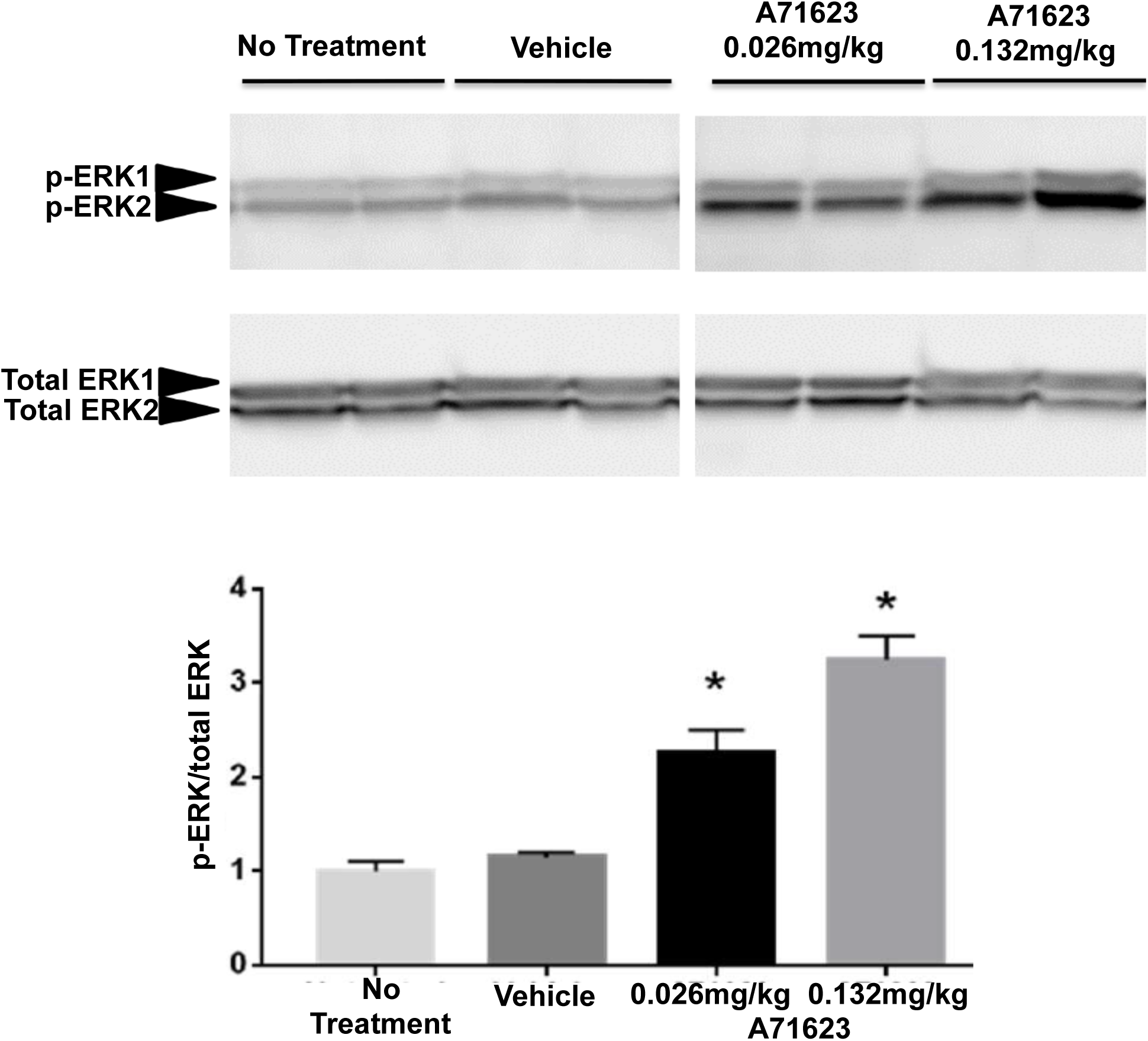
IP Injection of Cck1R agonist A71623 Increases Cerebellar Phospho-ERK1/2. A) Western blot of phosphor-ERK1/2 (P-ERK) and total ERK1/2. Two doses were administered i.p. to WT/FVB mice. Cerebella were harvested 24 hours post-injection. B) Quantification of (A). n=2 per group. One-Way ANOVA, tukey post-hoc test, *p<0.05.

**Figure S3.**
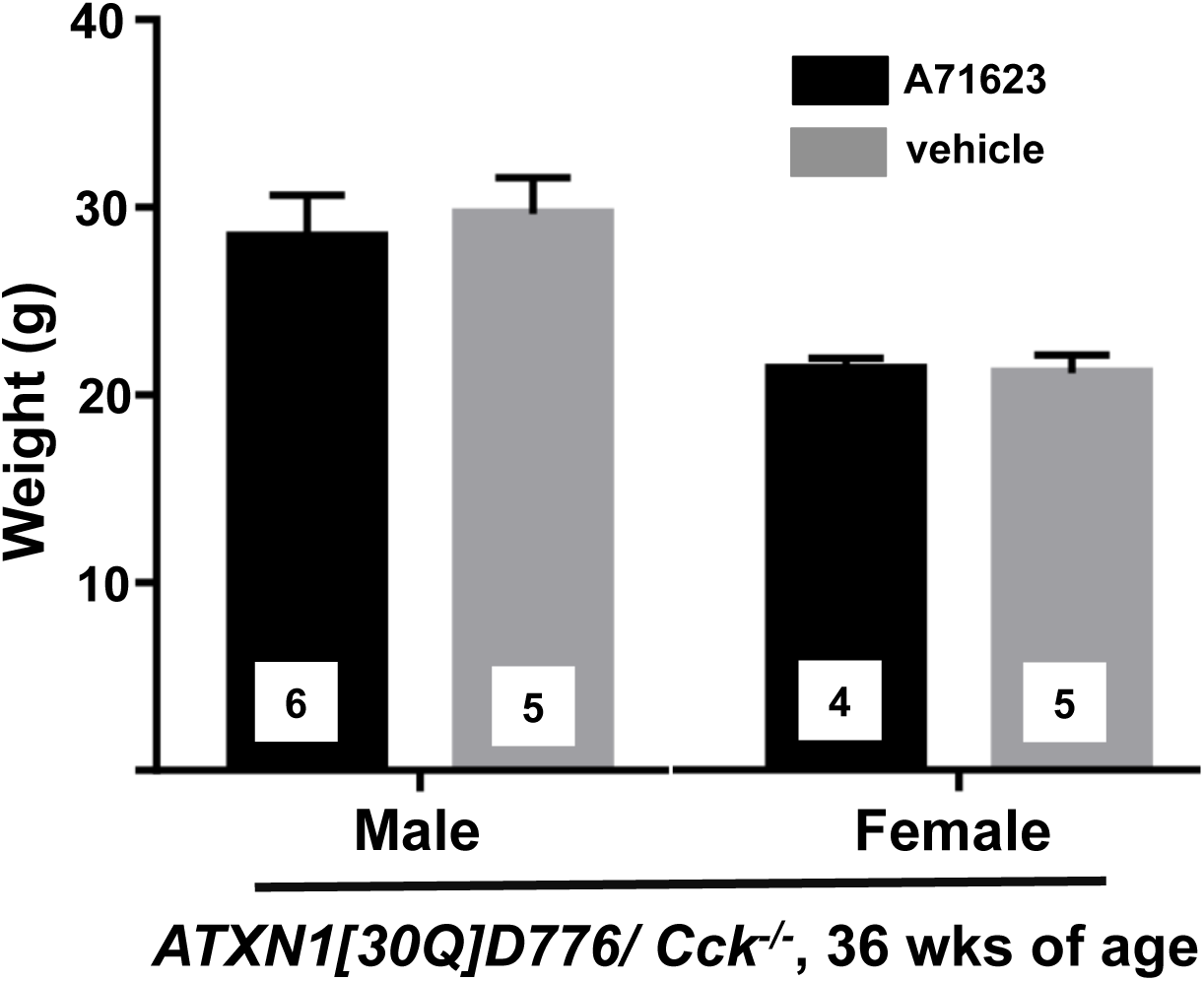
Administration of Cck1R agonist A71623 Does not Alter Weight of *ATXN1[30Q]-D776/Cck^-/-^* Mice. Male and female *ATXN1[30Q]-D776/Cck^-/-^* mice weighed following IP administration of A71623 (0.026mg, 30nmoles/Kg/day) or vehicle via osmotic pumps for 36 weeks.

**Figure S4.**
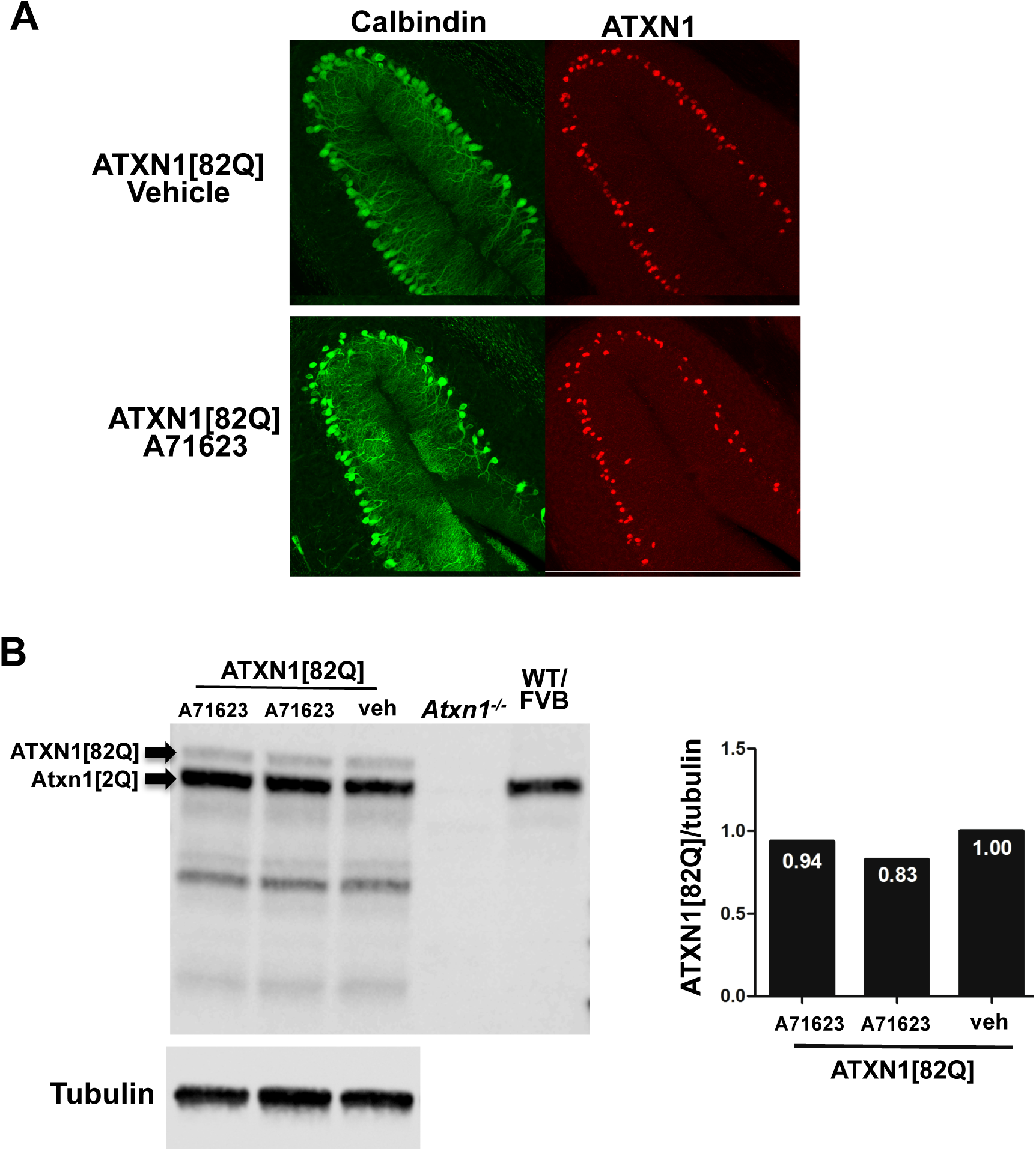
Administration of Cck1R agonist A71623 Does not Effect Expression of ATXN1[82Q] protein in Cerebella of *ATXN1[82Q]* Mice. A71623 (0.026mg, 30nmoles/Kg/day) or vehicle was administered for 12 weeks via IP osmotic pumps. A) Immunohistochemical staining of calbindin (Purkinje cell specific marker) and ATXN1 in Purkinje Cells. B) Western blot analysis of cerebellar ATXN1 expression.

**Figure S5.**
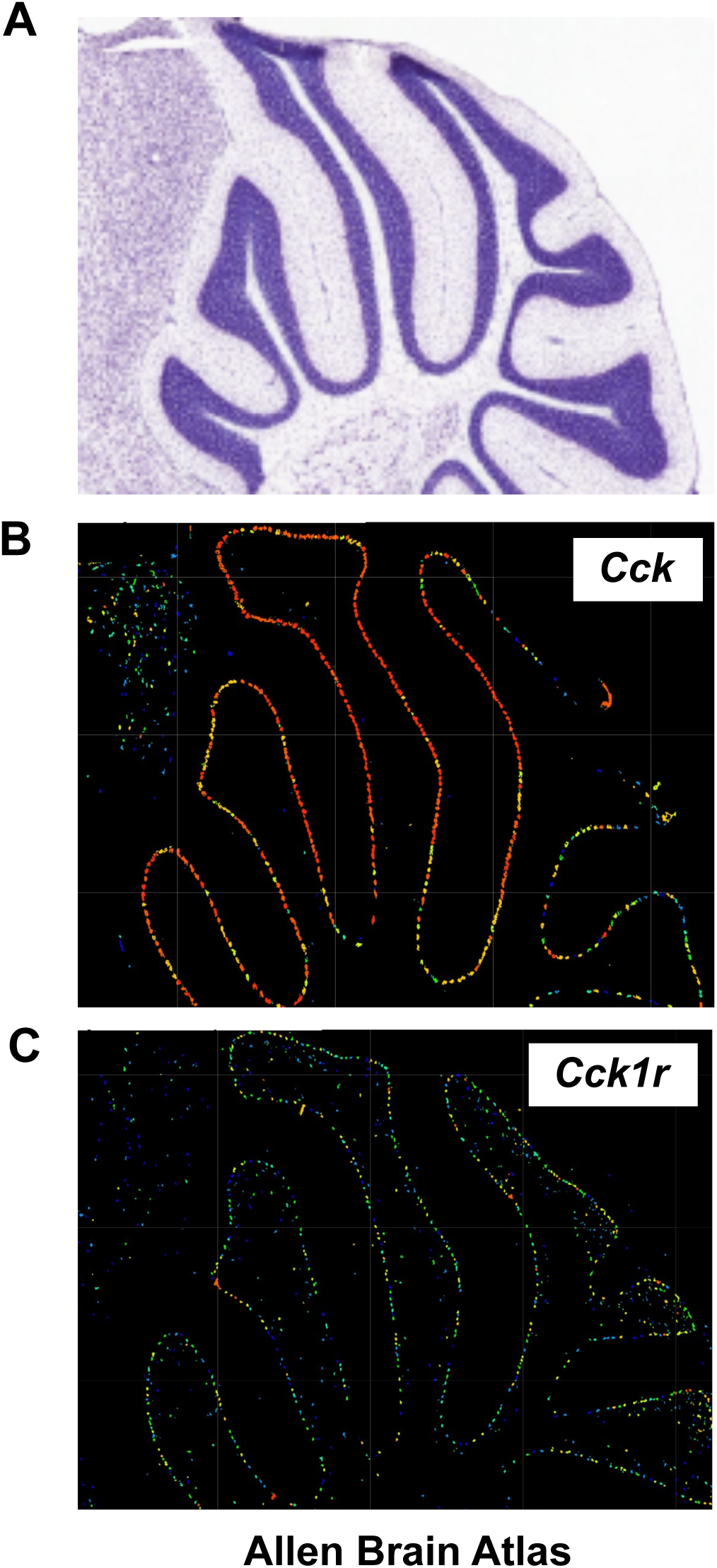
Purkinje cells Express *Cck* and *Cck1r*. A) Nissel stained sagittal vermal section. B) In situ hybridization for *Cck* RNA expression on a sagittal cerebellar vermal section. C) In situ hybridization for *Cck1***r** RNA expression on a sagittal cerebellar vermal section. All images are from Allen Brain Atlas; http://www.brain-map.org.

## REFERENCES

1. Angliker, N., Burri, M., Zaichuk, M., Fritschy, J.-M., and Rüegg, M.A. (2015) mTORC1 and mTORC2 have largely distinct functions in Purkinje cells. Eur. J. Neurosci. 42:2595–2612.

2. Asin, K.E., Bednarz, L., Nikkel, A.L., Gore, P.A., Montana, W.E., Cullen, M.J., Shiosaki, K., Craig, R., and Nadzan, A.M. (1992a) Behavioral effects of A71623, a highly selective CCK-A agonist tetrapeptide. Am. J. Physiol. 263:R125–R135.

3. Asin, K.E., Bednarz, L., Nikkel, A.L., Gore, P.A., and Nadzan, A.M. (1992b) A-71623, a selective CCK-A receptor agonist, suppresses food intake in the mouse, dog, and monkey. Pharmacol. Biochem. Behav. 42:699–704.

4. Burright, E.N., Clark, H.B., Servadio, A., Matilla, T., Feddersen, R.M., Yunis, W.S., Duvick, L.A., Zoghbi, H.Y. and Orr, H.T. (1995) SCA1 transgenic mice: a model for neurodegeneration caused by an expanded CAG trinucleotide repeat. Cell 82:937–948.

5. Cawson, E.E. and Miller, L.J. (2010) Therapeutic potential for novel drugs targeting the type 1 cholecystokinin receptor. British J. Pharmacol. 159: 1009–1021.

6. Clark, H.B., Burright, E.N., Yunis, W.S., Larson, S., Wilcox, C., Hartman, B., Matilla, A., Zoghbi, H.Y., and Orr, H.T. (1997) Purkinje cell expression of a mutant allele of *SCA1* in transgenic mice leads to disparate effects on motor behaviors followed by a progressive cerebellar dysfunction and histological alterations. J. Neurosci.17:7385–7395.

7. Dufresne, M., Seva, C., and Fourmy, D. (2006) Cholecystokinin and gastrin receptors. Physiol. Rev. 86:805–847.

8. Durr, A. (2010) Autosomal dominant cerebellar ataxias: Polyglutamine expansions and beyond. Lancet Neurol. 9:885–894.

9. Duvick, L., Barnes, J., Ebner, B., Agrawal, S., Andresen, M., Lim, J., Giesler, G.J., Zoghbi, H.Y., and Orr, H.T. (2010) SCA1-like disease in mice expressing wild type ataxin-1 with a serine to aspartic acid replacement at residue 776. Neuron 67:929–935.

10. Emamian, E.S., Kaytor, M.D., Duvick, L.A., Zu, T., Susan K. Tousey. S.K., Zoghbi, H.Y., Clark, H.B., and Orr, H.T. (2003) Serine 776 of ataxin-1 is critical for polyglutamine-induced disease in *SCA1* transgenic mice. Neuron 38:375–387.

11. Hansen, S.T., Meera, P., Otis, T.S., and Pulst, S.M. (2013) Changes in Purkinje cell firing and gene expression precede behavioral pathology in a mouse model of SCA2. Hum. Mol. Genet. 22: 271–283.

12. Ingram, M., Wozniak, E.A.L, Duvick, L., Yang, R., Bergmann, P., Carson, R., O’Callaghan, B., Zoghbi, H.Y., Henzler, C., and Orr, H.T. (2016) Cerebellar transcriptome profiles of *ATXN1* transgenic mice reveal SCA1 disease progression and protection pathways. Neuron 89:1194–1207.

13. Klockgether, T. (2011) Update on degenerative ataxias. Curr. Opin. Neurol. 24:339–345.

14. Koeppen, A.H. (2005) The pathogenesis of spinocerebellar ataxia. Cerebellum 4:62–73.

15. Kopin, A.S., Mathes, W.F, McBride, E.W., Nguyesn, M., Al-Haider, W., Schmitz, F., Bonner-Weir, S., Kanarek, R., and Beinborn, M. (1999) The cholecystokinin-A receptor mediates inhibition of food intake yet is not essential for the maintenance of body weight. J. Clin. Invest. 3:383–391.

16. Langfelder, P., and Horvath, S. (2008) WGCNA: an R package for weighted correlation network analysis. BMC Bioinformatics 2008, 9:559.

17. Lee, S.Y., and Soltesz, I. (2011) Cholecystokinin: a multi-functional molecular switch of neuronal circuits. Dev. Neurobiol. 71:83–91.

18. Miller, L.J. and Desai, A. J, (2016) Metabolic actions of the type 1 cholecystokinin receptor: Its potential as a therapeutic target. Trends Endocrinol, Metab., in press.

19. Orr, H.T., Chung, M.Y., Banfi, S., Kwiatkowski, T.J., Jr., Servadio, A., Beaudet, A.L., McCall, A.E., Duvick, L.A., Ranum, L.P. and Zoghbi, H.Y. (1993) Expansion of an unstable trinucleotide CAG repeat in spinocerebellar ataxia type 1. Nat. Genet. 4:221–226.

20. Paul, S., Dansithong, W., Figueroa, K.P., MS, Gandelman, M., Scoles, D.R. and Pulst, S.M. (2021) Staufen1 in human neurodegeneration. Ann Neurol, in press, DOI: 10.1002/ana.26069

21. Paulson, H.L., Shakkottai, V.G., Clark, H.B., and Orr, H.T. (2017) Polyglutamine spinocerebellar ataxias – from gene identification to potential treatments, Nature Revs. Neuroscience, 18: 613–626.

22. Robinson K.J., Watchon, M., and Laird, A.S. (2020) Aberrant cerebellar circuitry in the spinocerebellar ataxias. Front. Neurosci. 14:707.

23. Ruegseegger, C., Stucki, D.M., Steiner, S., Angliker, N., Radecke, J., Keller, E., Zubeer, B., Rüegg, M.A. and Saxena, S. (2016) Impaired mTORC1-dependent expression of Homer 3 influences SCA1 pathophysiology. Neuron 89: 129–146.

24. Seidel, K., Siswanto, S., Brunt, E.R.P., den Dunnen, W., Korf, H.-K., and Rub, U. (2012) Brain pathology of spinocerebellar ataxias. Acta Neuropathol. 124:1–21.

25. Su, K.-H. and Dai, C. (2017) mTORC1 senses stresses: coupling stress to proteostasis. Bioassays 39: 5, 1600268.

26. Watase, K., Weeber, E.J., Xu, B., Antalffy, B., Yuva-Paylor, L., Hashimoto, K., Kano, M., Atkinson, D.L., Sweatt, J.D., Orr, H.T., Paylor, R., and Zoghbi, H.Y. (2002) A long CAG repeat in the mouse Sca1 locus replicates SCA1 features and reveals the impact of protein solubility on selective neurodegeneration. Neuron 34:905–919.

27. Zu, T, Duvick, L.A., Kaytor, M.D., Berlinger, M., Zoghbi, H.Y., Clark, H.B. and Orr, H.T. (2004) Recovery from polyglutamine-Induced neurodegeneration in conditional *SCA1* transgenic mice. J. Neuroscience 24:8853–8861.

